# Hamster and ferret experimental infection with intranasal low dose of a single strain of SARS-CoV-2

**DOI:** 10.1101/2020.09.24.311977

**Authors:** Elodie Monchatre-Leroy, Sandrine Lesellier, Marine Wasniewski, Evelyne Picard-Meyer, Céline Richomme, Franck Boué, Sandra Lacôte, Séverine Murri, Coralie Pulido, Johann Vulin, Francisco J Salguero, Meriadeg Ar Gouilh, Alexandre Servat, Philippe Marianneau

## Abstract

Understanding the pathogenesis of the SARS-CoV-2 infection is key to develop preventive and therapeutic strategies against COVID-19, in the case of severe illness but also when the disease is mild. The use of appropriate experimental animal models remains central in the in-vivo exploration of the physiopathology of infection and antiviral strategies. This study describes SARS-CoV-2 intra-nasal infection in ferrets and hamsters with low doses of low-passage SARS-CoV-2 clinical French isolate UCN19, describing infection levels, excretion, immune responses and pathological patterns in both animal species. Individual infection with 10^3^ pfu SARS-CoV-2 induced a more severe disease in hamsters than in ferrets. Viral RNA was detected in the lungs of hamsters but not of ferrets and in the brain (olfactive and/or spinal bulbs) of both species. Overall, the clinical disease remained mild, with serological responses detected from 7 days and 10 days post inoculation in hamsters and ferrets respectively. Virus became undetectable and pathology resolved within 14 days. The kinetics and levels of infection can be used in ferrets and hamsters as experimental models for understanding the pathogenicity of SARS-CoV-2, and testing the protective effect of drugs.

## INTRODUCTION

Coronaviruses Disease 2019 (COVID-19) is induced by the Severe Acute Respiratory Syndrome coronavirus 2 (SARS-CoV-2) (1)] and has become a pandemic causing extensive economic, social, medical and scientific disruption around the world. Since 2002, SARS-CoV-2 is the third epidemic coronavirus transmitted from animals to humans after SARS-CoV (2)] and Middle East Respiratory Syndrome coronavirus (MERS-CoV) (3)] and the magnitude of the COVID-19 in term of the number of human cases is by far the greatest. To fight COVID-19, neither vaccines nor therapeutic treatments are currently available. Understanding the pathogenesis of the SARS-CoV-2 infection is key to maximize prevention and to develop therapeutic solutions, and this is possible through susceptible animal models such as non-human primates (cynomolgus (*Macacca fascicularis*) and rhesus (*Macacca mulatta*) macaques), cats, hamsters and ferrets (4–13)], being the two latest species the most economic and easiest to house and handle. To our knowledge, all ferrets and hamsters experimental infection studies (9,13),] published so far, apart from two studies (9,13),] were performed with high doses of SARS-CoV-2 (ranging from 8.10^4^ TCID50 to 10^5.5^ TCID50 or 10^6^ pfu per animal), independently of animal weight and mainly with the objective of inducing severe infections. Here, we choose to explore the effects of a lower viral infection dose, considered closer to natural and more common infection conditions with SARS-CoV-2 in humans, probably leading to mild disease symptoms, but with potential viral excretion.

The main objectives of the study were a) to characterize the kinetics of the disease (clinical signs, pathogenicity and immune responses) in ferrets and hamsters infected with low doses of low-passage SARS-CoV-2 clinical isolate, in order to develop suitable animal models for therapeutic and vaccine studies.

## METHODS

### 1. SARS-CoV-2 virus

SARS-CoV-2 viral strain UCN19 was isolated during the course of the active epidemic from naso-pharyngeal flocked swabs obtained from patients at the University Hospital of Caen, Normandy, France, who were suffering from respiratory infection and were confirmed infected by SARS-CoV-2 by routine molecular diagnosis. The swabs were eluted in UTM media (Copan, Italy) at 4°C for less than 48 hours. Vero CCL-81 cells (passage 32, from ATCC, USA), grown at 80% confluence level were inoculated with 200µl micro-filtered elution. Cells were visually checked for cytopathic effect on a daily basis using an inverted microscope. Cell supernatants (12 ml) were harvested at day 3 after inoculation and immediately used for passage 1 (P1) produced in T75 culture flasks containing Vero cells as previously described. P1 was used for stock production of UCN19, aliquoted and stored at −80°C before titration, genomic quantification, sequencing and experimental infections to ferrets and hamsters. The use of the low-passage SARS-CoV-2 clinical isolate P1 reduces the risk of cell-culture induced genetic modification.

### 2. Animal experimental design

The experimental protocols complied with the regulation 2010/63/CE of the European Parliament and of the council of 22 September 2010 on the protection of animals used for scientific purposes (14)] and as transposed into French law (15)]. These experiments were approved by the Anses/ENVA/UPEC ethic committee and the French Ministry of Research (Apafis n°24818-2020032710416319).

Fifteen 10 month-old ferrets (*Mustela putorius furo*, ten neutered males and five females Euroferrets, Denmark) and twenty-one 8-week old female hamsters (*Mesocricetus auratus*, strain RjHan:AURA - Janvier Labs, France) were used. Ferrets and hamsters were kept in cages with environmental enrichment, allocated into groups of 2 to 5 ferrets and 2 hamsters per cages, with non-infected animals kept in separate room from the infected animals. Food and water were provided ad-libitum. Weight, body temperature (measured once a day by subcutaneous chips IPTT300, Plexx, the Netherlands) and activity levels of all animals were monitored and recorded on a daily basis throughout the duration of the experimental procedures.

Twelve ferrets and fifteen hamsters were anesthetized with isofluorane and inoculated by the intranasal route with 2.10^3^ pfu and 1.8.10^3^ pfu of UCN19 SARS-CoV-2 strain respectively. The doses ranged from 1 to 3 pfu per gram of ferret (average 2 pfu/g ferret) and from 20 to 25 pfu per gram of hamster (average of 22 pfu/g of hamster). Ferrets received the inoculum in 250 µl and hamsters in 20 µl in each nostril. At day 2 (D2) post-inoculation, D4, D7, D10 and D14, infected animals were anaesthetized with isoflurane for sample collection: oro-pharyngeal swabs (plain swab rayon tipped, Copan, Italy), nasal washes in ferrets only (by administering 500ml PBS to each nostril) hamsters being too small for this sampling technique, rectal swabs in ferrets (plain swab rayon tipped, Copan, Italy) and fecal pellets spontaneously produced in hamsters. On the same time-points, blood was collected from the cranial vena cava in ferrets using vacutainer (SST tubes, BD) and from the retro-orbital vein in hamsters to isolate serum for immunological tests. Hamster serum was tested by RT-PCR and viral culture. The number of animals sampled at each time-point decreased overtime with six inoculated animals (three ferrets, including one female ferret and three hamsters) euthanized sequentially at D2, D4, D7, D10 (hamsters only) and D14 and submitted to post-mortem (PM) examination. At PM, the following samples were collected and stored frozen at −80 °C for viral RNA measurement and viral culture or in 10% buffered formalin for histology: nasal turbinates, tonsils (ferrets only / tissue absent in hamster’s anatomy), proximal trachea, lung lobes (left lobes for RNA extraction and viral titration and right lobes for histopathology), liver, spleen, large intestine, kidneys, olfactory bulb and spinal bulb. In addition, broncho-alveolar lavage (collected with PBS in ferrets only for practical reasons) and urine were collected at PM. Three ferrets and six hamsters were inoculated with PBS as naive controls and were sampled at the end of the study to confirm their negative status and provide background data for immunological studies.

### 3. Virus titration

Viral load was determined for viral inoculum and for a subset of samples (list in the supplementary table) by plaque assay on VeroE6 cells. A 12 well-plate was seeded with a VeroE6 cell suspension 24 hours before virus inoculation. Organs were harvested, weighted and homogenized in appropriate volumes of DMEM with stainless steel beads (QIAGEN) for 3min at 30 Hz using TissueLyserII (QIAGEN). Homogenates were then clarified by centrifugation (2,000 × g at 4°C for 10 min), aliquoted and stored at −80°C. A tenfold serial dilution of the samples (clinical samples and tissue homogenates) was performed in DMEM supplemented with FCS and penicillin/streptomycin. Inoculum was added in each well and the plate was incubated at 37°C in 5% C02 for 1 hour. At least one uninfected well was used as an independent negative control. One hour later, plaque assays were overlaid with carboxy-methylcellulose mix (CMC 3.2% DMEM 5% FBS (V/V)) and after 5 days were fixed and stained with a crystal violet solution (3.7% formaldehyde, 0.2% crystal violet). The titer in PFU/mL (number of plaques / (dilution x volume of diluted virus added to the well) was determined by dividing the number of plaques for the adequate dilution by the total dilution factor.

### 4. RNA extraction and TaqMan RT-qPCR

Tissue homogenates and clinical samples were treated the same way from that point. The adequate volume of Triton X-100 (MP Biomedicals, Illkirch, France) was added to AVL Lysis buffer (Qiagen, Courtaboeuf, France) to reach a concentration of 2.7 %, in order to inactivate potential infectious status of samples by SARS-CoV-2. A negative control RNA extraction was performed for each set of 12 samples tested. TaqMan RT-qPCR was performed with the SuperScript™III Platinum™One-Step qRT-PCR kit (Invitrogen, Fischer Scientific, Illkirch, France) using the protocol described by Corman et al. (16)]. Coronavirus primers (E_Sarbeco_F and E_Sarbeco_R) and probe (E_Sarbeco_P1 labelled with the fluorescent dye FAM-BHQ1) targeting the envelope protein gene (E gene) were used for this study. Primers and probe provided by Eurogentec (Angers, France). All TaqMan RT-qPCR assays were performed on the thermocycler Rotor Gene Q MDx (Qiagen, Courtaboeuf, France) and Lightcycler LC480 (Roche, France). Negative and positive controls were included in each RT-qPCR assay. Positive controls were performed for each assay by testing in duplicates 5 serial 10-fold dilutions of a calibrated SARS-CoV-2 RNA titrating 3.10^6^ copies/µL of RNA. The determination of SARS-CoV-2 RNA titer in number of copies/µL was determined by testing a standard curve with six 10-fold dilutions of a SARS-CoV-2 RNA titrating 3.10^6^ E gene copies/µL of RNA extracted from the SARS-CoV2 strain UCN19 and itself quantified using an E gene transcript. A threshold setting (Ct) of 0.05 was used as the reference for each RT-qPCR assay. The efficiency, slope and correlation coefficient (R^2^) were determined with the Rotor Gene software. All reactions were carried out as technical duplicates. A cut-off > 35 was defined for low positive results (>300 copies/µL of RNA)

### 5. Histological analyses

Samples from the right cranial and caudal lung lobes together with tonsil (in ferrets only) and trachea were fixed by immersion in 10% neutral-buffered formalin and processed routinely into paraffin wax. Four µm sections were cut and stained with haematoxylin and eosin (H&E) and examined microscopically. In addition, samples were stained using the RNAscope in situ hybridization (ISH) technique to identify the SARS-CoV-2 virus RNA as previously described (9). Briefly, tissues were pre-treated with hydrogen peroxide for 10 mins (RT), target retrieval for 30 mins (98-101°C) and protease plus for 30 mins (40°C) (Advanced Cell Diagnostics). A V-nCoV2019-S probe (Advanced Cell Diagnostics, Biotechne) was incubated on the tissues for 2 hours at 40°C. Amplification of the signal was carried out following the RNAscope protocol using the RNAscope 2.5 HD Detection kit – Red (Advanced Cell Diagnostics, Biotechne).

### 6. Serological analyses

The serum of six different animals (three ferrets and three hamsters) were sampled at each time-point, except at D10 and D14 when the same ferrets were tested consecutively. The plates were coated with SARS-CoV-2-infected Vero E6 cell lysates at 4°C overnight. Non-infected Vero E6 lysate was also coated to provide sample background OD values. These antigens were previously validated with negative and positive human sera. Serum samples, diluted 1:100, were incubated for 1 h at 37°C. Specific antibody binding was detected by peroxydase-labelled Goat anti-Hamster IgG (H+L) (Invitrogen) and peroxydase-labelled Goat anti-Ferret IgG (H+L) (KPL), diluted 1:5000 and 1:100 respectively. Substrate (TMB single solution, Fisher Scientific) was added after 1h at 37°C and the reaction was stopped with 10.6% H3PO4. The optical density was determined on both viral and negative control at 450nm for both species. Net OD values were calculated by removing the sample background OD value from OD sample values.

The presence of neutralizing antibodies was detected by performing a plaque reduction neutralization test (PRNT) using P2 of SARS-CoV-2 UCN19 strain. Hamster and ferret sera were serially diluted and mixed with an equal volume containing 100 plaque forming units (pfu) of virus per 125µl. The mixtures were incubated for 1h and subsequently inoculated into wells of twelve-well tissue plates containing confluent Vero E6 cells. After adsorption for 1h, the wells were overlaid with a mixture of CMC (carboxymethylcellulose sodium salt low viscosity) and 5% FCS DMEM’s medium. Plates were incubated for five days and finally colored by violet Crystal. A 50% reduction in the number of plaque was used as the criterion for virus neutralization titers. Endpoint titers of neutralizing antibodies were determined.

### 7. Statistical analyses

Wilcoxon signed-rank test was performed by package stat in R (version 3.3.3) to compare viral RNA quantities in nasal washes between males and females ferrets and with GraphPad Prism v6 to compare the areas under the curves for the kinetics of viral presence in clinical samples and of weight changes from D0. Two ways ANOVA was calculated (significance level fixed at 0.05, to analyse the effect of “species” and “time” on RNA levels recovered in tissue samples collected from ferrets and hamsters using GraphPad Prism v6.

## RESULTS

### 1. Clinical observations

The experimental SARS-CoV-2 infection did not result in any death and no clinical endpoint was reached in any animal. Lethargy was observed on days 7 and 8 post infection in three ferrets. Snoring was also reported in one ferret between D7 and D14. No hyperthermia was observed in any infected animal at any point compared with controls. Weight remained stable in ferrets. In hamsters, an average weight gain was observed in animals overtime, in average reduced, although not significantly, in the inoculated group (Figure 1).

**Figure 1.**
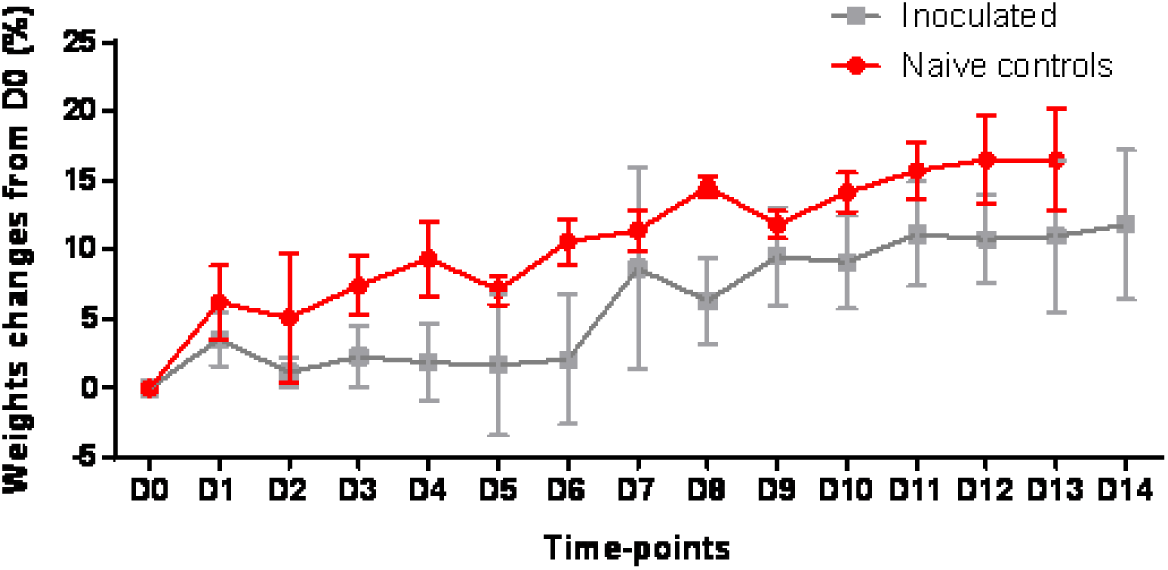
Hamsters weight changes from D0 (%) with mean and standard deviation after inoculation with SARS-CoV-2 or PBS (in the naïve controls).

### 2. Local viral RNA presence measured by RT-PCR

Viral RNA was recovered in a large range of tissues in both species from two days post infection and tended to decrease over time, with almost all tissues negative by D14, except in 1/ in ferrets, nasal turbinates (Figure 2A), tonsils (tissue absent in hamster’s anatomy) (Figure 2B), spinal bulb (Figure 2J) and 2/ in hamsters, the trachea (Figure 2C) and the lungs (Figure 2D). More variability was observed in RNA levels within ferret groups than hamster groups at each time-point. The most heavily infected tissues were 1/ the nasal turbinates with no significant difference between average infection levels measured in ferrets and hamsters, 2/ the trachea significantly more infected in hamsters than in ferrets and 3/ the lungs with high RNA levels in hamsters and no detection in ferrets. Viral RNA was recovered at lower levels than in the respiratory tissues in the liver (Figure 2E), the spleen (Figure 2F) and the kidneys (Figure 2H) on D2 and D4 in hamsters only. In the large intestine (Figure 2G), the average viral RNA levels were similar in ferret and hamster groups, although only individual ferrets presented values higher than hamsters at D2, D4 and D7. In the olfactive bulb (Figure 2I), viral RNA was recovered in both species, with higher average levels in hamsters at D2 and D4, but a longer persistence in ferrets (up to D14). All three hamsters were positive in the spinal bulb at D2 but subsequently became negative, while only one ferret was positive at D7. Urine samples were negative at all time-points in both species, as well as retro-pharyngeal lymph nodes (tested in ferrets only) and hamster serum (data shown in the complementary table S1).

**Figure 2.**
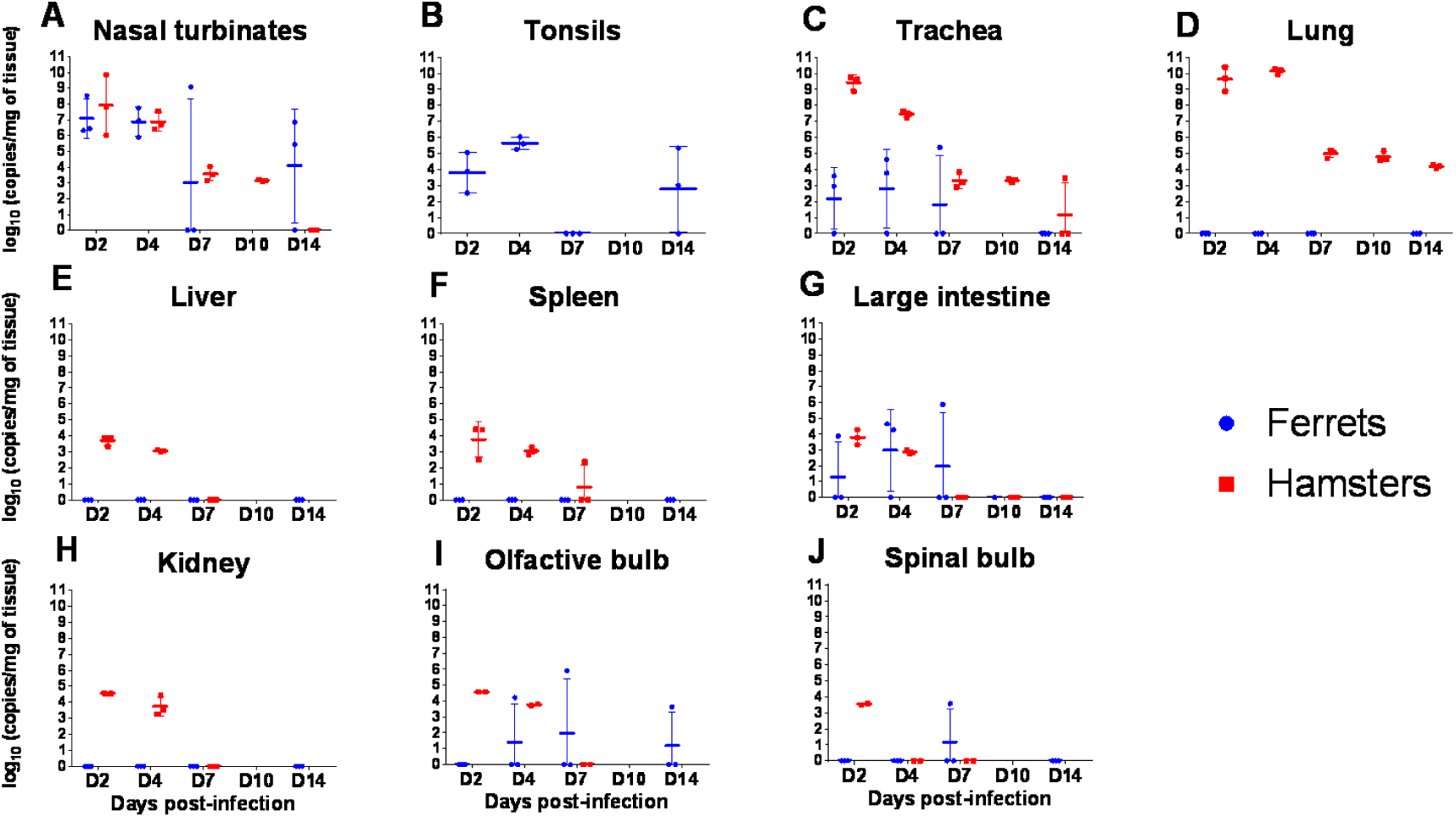
Viral RNA loads in ferrets and hamsters inoculated with UCN19 SARS-CoV-2.

Viral RNA was found in oral swabs of both species from D2 to D10 (Figure 3A) with no oral swab positive at D14 in any species. The areas under the curves did not differ between hamsters and ferrets. Nasal wash fluids were positive from D2 to D14 (Figure 3B) (collected in ferrets only) and this result was consistent with high levels of viral RNA found in nasal turbinates in this species (Figure 2A). The viral RNA levels in nasal washes were not different between males and females ferrets independently from the experimental time point (p-value of 0.28 on D2, 0.55 on D4 and of 1 on D7, D10 and D14). RNA levels decreased faster in rectal swabs (Figure 3C) than in the other clinical samples and were all negative by D10, with no difference between areas under the curves of ferrets and hamsters.

**Figure 3.**
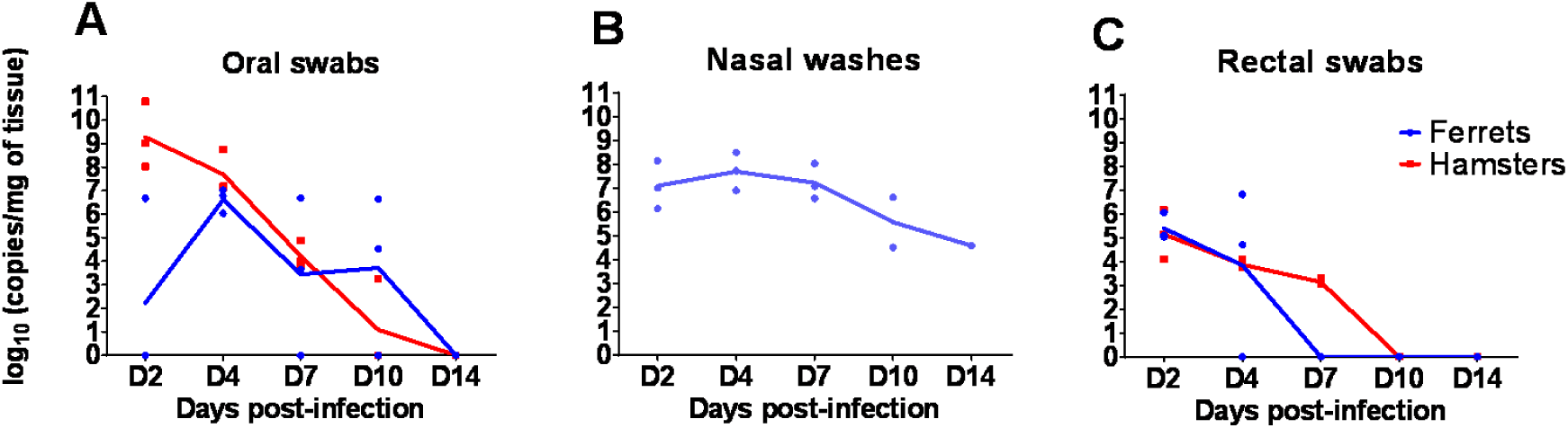
Viral RNA loads in clinical samples. (A): oral swabs, (B): nasal washes, (C): rectal swabs collected from to D2 to D14 in three ferrets and three hamsters inoculated intra-nasally with UCN19 SARS-CoV-2 (nasal washes were only collected in ferrets).

### 3. Virus titration

In hamsters, infectious virus was detected from nasal turbinates, trachea and lungs in all the necropsied hamsters at D2 and D4 with 2.7.10^0^ to 1.32.10^4^ pfu/ml tissue and from oral swabs for two out of three hamsters at D2 (4.10^1^ and 2.4.10^2^ pfu/ml) and one at D4 (8.0.10^1^ pfu/ml). In ferrets, infectious virus was detected in turbinates and nasal washes, but not in the lungs. Nasal turbinates were found infectious (in four out of nine samples) between D2 and D7 with a maximal titer of 1.28.10^4^ pfu/ml at D7. Almost all nasal washes (five out six samples) were found infectious between D2 and D4 with a maximal titer of 3.04. 10^3^ pfu/ml at D2.

No infectious virus was isolated in any other tested sample (see supplementary data in tables S1 and table S2).

### 4. Histopathology

In hamsters, inflammatory infiltration was observed in the lung from D2 to D7, with presence of macrophages, lymphocytes and neutrophils, usually surrounding airways and within the alveolar walls (Figure 4). These inflammatory infiltrates showed signs of cell death and high amount of viral RNA detected by ISH, mostly at D2 and D4 (Figure 4), and at much lower level at D7. Animals culled at D10 and D14 did not show any remarkable changes in the lungs and no presence of viral RNA. In the trachea, mild inflammatory infiltration of the epithelium was observed together with minimal to mild epithelial necrosis at D2, D4 and D7. Viral RNA was very abundant within the tracheal epithelial cells at D2, and decreased from D4 with a complete absence from D7 onwards (Figure 4)

**Figure 4.**
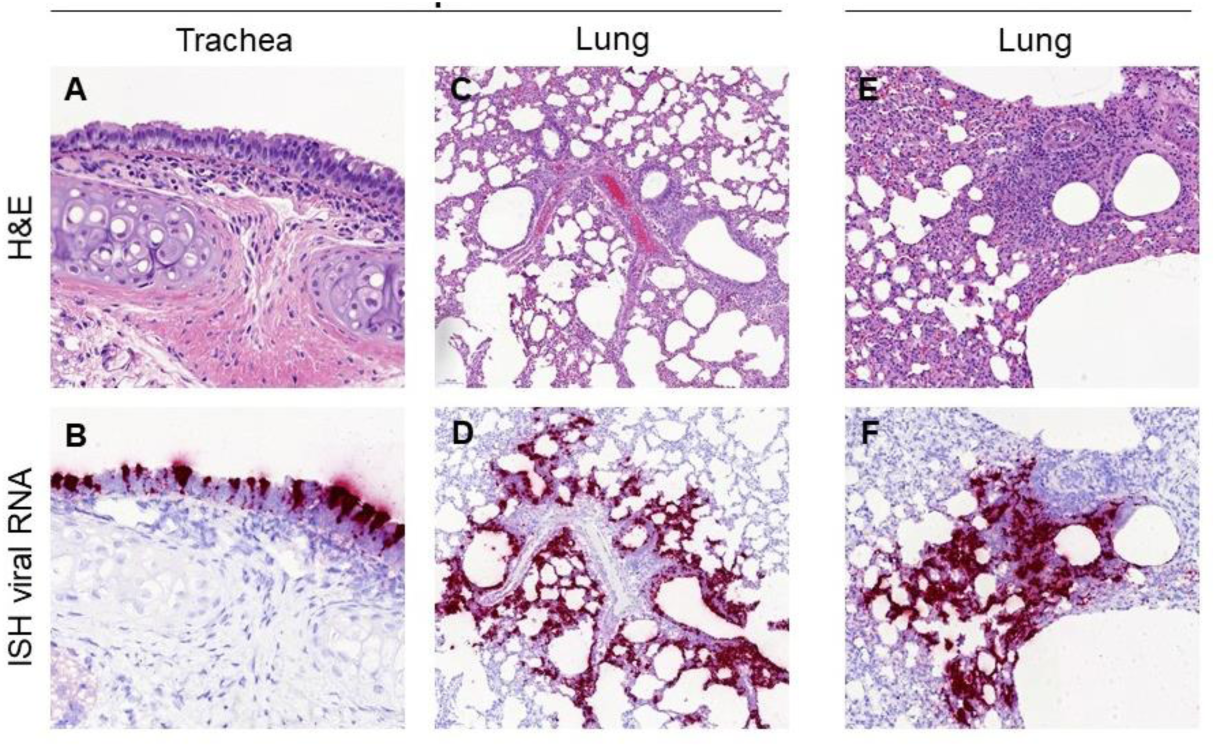
Histopathological findings in hamsters inoculated with SARS-CoV-2 UCN19 strain. Mild inflammatory cell infiltration observed in the trachea at D2 (A, 400x) with high presence of viral RNA by ISH within respiratory epithelial cells (B, 400x). Inflammatory infiltrates with the lung parenchyma, mostly within the bronchial and bronchiolar mucosa but also surrounding airways and blood vessels are observed at D2 (C, 100x) and D4 (E, 200x). The presence of the inflammatory infiltrates is correlated with the viral RNA staining in sequential sections at D2 (D, 100x) and D4 (F, 200x).

In ferrets, histopathological findings were less severe than in hamsters. Non-inoculated ferrets showed only minimal to mild inflammatory cell infiltrates around the bronchioles and blood vessels (peribronchiolar and perivascular cuffing). In inoculated animals, mild bronchiolitis was observed at D2 in all animals with presence of inflammatory cell infiltrates within the bronchiolar luminae, mostly neutrophils but also some eosinophils, macrophages and lymphocytes. Perivascular and peribronchiolar cuffing was also observed (mainly mononuclear cells) (Figure 5). One of the three animals culled at D2 showed very few scattered cells positive to viral RNA within alveolar walls, not related to any histopathological lesion (Figure 5B). At D4, D7 and D14, individual animals showed mild to moderate perivascular cuffing, together with areas of bronchiolitis similar to those seen in animals culled at D2 (Figure 5). Small areas of consolidation (inflammatory infiltrates composed mainly of macrophages and lymphocytes within the parenchyma) were also observed (Figure 5F). No remarkable changes were observed in the tonsil nor the proximal trachea, with only few scattered positive cells to viral RNA (ISH) in the trachea at D2 not related to any histopathological lesion.

**Figure 5.**
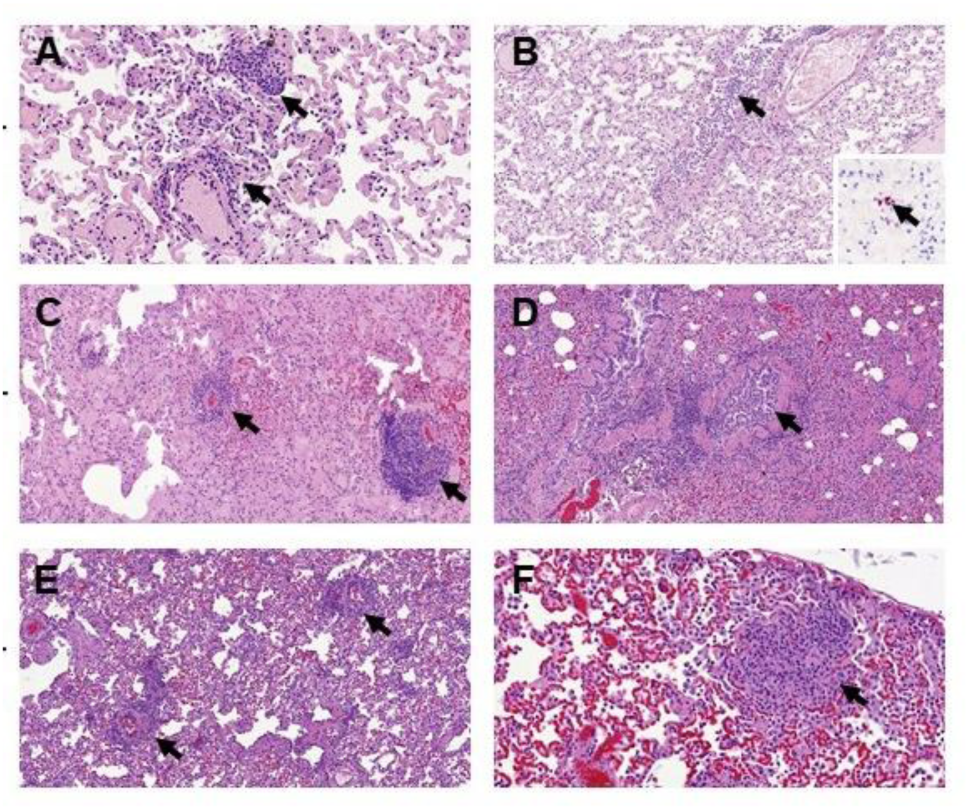
Histopathological findings in the lungs of ferrets inoculated with the UNC19 SARS-CoV-2 strain. Perivascular cuffing (A, 400x, arrows) and mild bronchiolitis (B, 200x, arrow) was observed at D2, with minimal presence of viral RNA (B, insert, 400x) within alveolar walls not related to histopathological lesions. Perivascular cuffing was also observed at D4 (C, 200x, arrows) and D7 (E, 200x, arrows). Mild bronchiolitis with presence of intraluminal inflammatory infiltrates was observed at D4 (D, 200x, arrow). Scattered foci of parenchymal inflammation were also observed at D7 (F, 400x, arrow)

### 5. Serology

A detectable level of IgG appeared from D7 in hamster serum and from D10 in ferrets. The level of IgG increased rapidly between D4 and D7 and remained consistently high until D14 in hamsters (Figure 6A), while in ferrets the increase appeared more gradual from D10. Neutralising antibodies were detected from D7 and D10 in hamsters and ferrets respectively (Figure 6B). The IgG levels did not correlate with the neutralizing titers. Indeed, in hamsters, the neutralizing titers halved between D7 and D14 whereas the level of detected IgG remained stable. More variability was observed in ferret responses at D10 and D14 than in hamsters. For one ferret, the level of neutralising antibodies was halved at D14 compared with D10, as observed for the hamsters. For another ferret, the response doubled from D10 to D14, while the last ferret of the group presented persistently the highest responses on D10 and D14 (Figure 6B).

**Figure 6:**
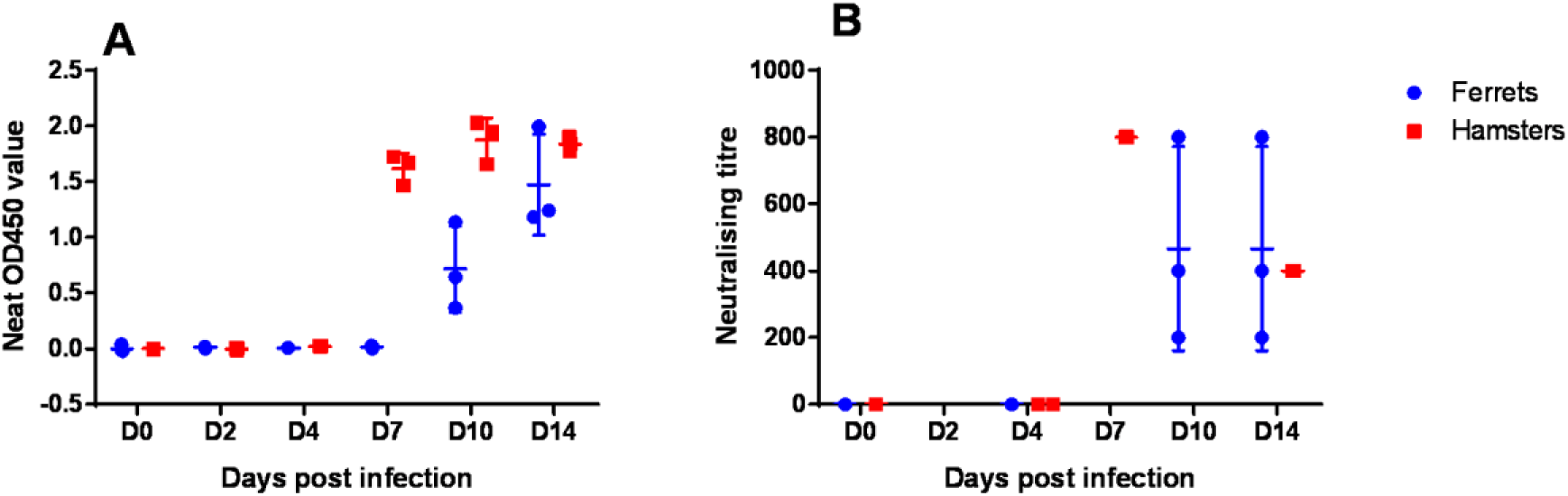
Antibody responses (mean and standard deviation) measured at each time-point in inoculated hamsters and ferrets A: Neat OD450 for IgG ELISA and B: Sero-neutralisation.

## DISCUSSION

A number of studies have already been published on the experimental infection of ferrets or hamsters with SARS-CoV-2, but none of them compared the two models with the same strain. Differences between protocols and strains may impact the clinical, immunological and pathological outcomes, and therefore impair the comparison of independent investigations. Moreover, most studies have focused on high inoculation doses in an attempt to induce severe forms of COVID-19 and understand their pathogenicity. In this study, we have focused on milder forms of COVID-19, more common than severe clinical course in human patients that also require preventive and therapeutic approaches. Only one other study (13) used a similarly low SARS-CoV-2 dose of 10^3^ pfu inoculated to young female hamsters, but it was delivered simultaneously by the intranasal and intraocular routes. Our study was planned to evaluate the pathogenicity and viral load, and peripheral immunogenicity in ferrets and hamsters inoculated by the intra-nasal route with a single low dose of the same SARS-CoV-2 clinical strain obtained after a single passage on cell culture. The infection of all inoculated hamsters and ferrets was confirmed in live animals and at post-mortem.

In experimentally infected hamsters, the clinical signs are generally considered limited, except for the significant weigh loss often reported. In our study, all hamsters tended to gain weight overtime although at different rates, in spite of local evidence of infection. Inoculated ferrets generally present even milder clinical signs of disease, independently from the challenge dose (8–11), without any weight loss, as reported in our study. The marked lethargy observed in ferrets in this study was also reported by others (8–11). It is possible that transient hyperthermia was missed if shorter than 24h (time interval between SC IPTT300 chips reading).

The kinetics, distribution and levels of RNA recovered in hamster tissues were similar to those reported in other studies (6,7,12), including with higher challenge doses, and generally higher than in ferrets, except in the nasal turbinates and large intestine where both species appeared to be on average infected at similar levels. The pathogenicity of SARS-CoV2 in the nasal cavity of the hamsters infected in this study involved respiratory and olfactory epithelium, including sustentacular cells (17). In ferrets, similar infection pattern has also been observed (9). In this species, nasal washes provided non-invasive evidence of local infection with consistently high level of viral RNA recovered from D2. The persistence of viral RNA over a long time-span increases the potential for detecting the infection, but should not be considered representative of presence of infectious virus. The infectiousness of nasal washes was demonstrated in VERO6 cells on D2 and D4 only, therefore on a short period of time and before the onset of clinical signs (lethargy) and the development of detectable serological responses. Such samples would be valuable to collect in hamsters in future experimental studies. In oral swabs, similar kinetics of viral RNA were observed in both species with a gradual decrease from D2 to complete absence on D14. The lack of positive sample in the retro-pharyngeal lymph nodes collected in ferrets suggests that SARS-CoV-2 did not drain through the oral mucosal, but targeted the nasal mucosa, infecting the trachea in both species, more severely in hamsters based on viral RNA recovery and histopathology findings, and finally the lungs, mostly in hamsters.

Both species developed a viral infection in the higher respiratory tract; however, hamsters developed a more severe infection in the lungs (measured by RNA levels and infectious virus isolation and histopathology) than ferrets. In ferrets, SARS-CoV-2 presence in lungs was only detected in very few cells in by RNAscope ISH technique, and not related to the histopathological lesions ISH was more sensitive than RNA extraction and cell culture in this study, detecting probably only small pieces of degraded viral RNA. This difference of sensitivity may also originate from the dilution effect of the virus in tissues, the heterogeneous viral distribution (although the infection protocols did not target any specific part of the lung), or the inhibitory effect of blood contained in samples on RNA detection. One study (13) in hamsters highlighted that lesions were more severe in right lobes than in left lung lobes; in our study, samples from the left lung lobes were used for RNA extraction and if this difference between lung sides also occurs in ferrets, it may have induced an under-estimation of infection in this species. The right lung lobes submitted for histopathology presented signs of viral infection. The impact of a species difference (between ferrets and hamsters) in pulmonary pathogenicity at the inoculation dose used in these species is unclear. SARS-CoV-2 is detected in the lungs of the majority of human fatal cases (18–20)] and ISARIC clinical investigations (21) reported shortness of breath or dyspnea within the five as most frequent symptoms, which is indicative of common viral invasion of the lungs by SARS-CoV-2. However lung viral analysis are not available in mild and moderate COVID-19 patients cases. The hamster model is a good proxy to study the invasion of the lungs by SARS-CoV2 and may therefore bear some resemblance with more serious human cases, in spite of showing limited clinical signs such as weight loss.

SARS-CoV-2 RNA was also measured in our study in the intestinal tract of ferrets and hamsters, although at a lower level than in the upper respiratory tract. As in most human cases (22), no infectious virus was isolated from intestinal samples, while viral RNA was detected. The presence of SARS-COV-2 in the brain has also been reported in other experimental studies (6,10,11), and rarely in human patients, although not systematically. In our study, viral RNA was recovered in both species mostly in the olfactive bulb but also in the spinal bulb. The impact of this dissemination in unclear.

Antibody kinetics in ferrets and hamsters were comparable with 1/ those seen in patients with a seroconversion between D7 and D14 (22), and 2/ those seen in experimental models using hamsters and ferrets (6,8–11). In our study, the increase in antibody levels occurred faster in hamsters than in ferrets, from no detectable level at D4 to consistently high levels between D7 and D14 in hamsters and between D7 and D10 in ferrets. These observations suggest that in these species, the exposure to low challenge viral dose inoculated by the intranasal route induced an early, significant and potentially protective humoral response. For both species, the rise of antibody levels slightly preceded or was concomitant with the disappearance of infectious virus (viral clearance) but further investigations would be required to conclude on the protective effect conferred by antibodies.

According to previous studies (9), reducing the challenge dose beyond the one used in our study impacts the reproducibility of infection success (only one out of six ferrets inoculated with 5.10^2^ pfu became infected in Ryan et al., experiments (9)). Infection doses should also be expressed and standardized per animal weight. In our study, we followed standard protocols by delivering homogeneous challenge doses between animals. As a result, the dose per weight was in the order of ten times higher in hamsters than in ferrets, which can contribute to the higher pathogenicity observed in this species. In addition, the females received a larger dose/kg than males since the same bolus was delivered to all animals in spite of different weights (average weight of 1419 g for males and of 740 g for females at D0). A lower susceptibility has been suggested in females, at least in humans (23–26), and this may explain why no difference in the severity of the disease was measurable between sexes in our study. Weight dependent infection doses in future studies may highlight sex differences in the severity of experimental outcomes.

COVID-19 is less frequently reported in children than in adults and elderly people (27)]. Two studies in hamsters (12,13) reported more severe lesions and clinical signs in older animals. The oldest ferrets used in experimental studies were 20 month old (8–11)], therefore no conclusion can be drawn on the effect of age in this species. Some studies suggest that thymus produced thymosin may have protective effect (28,29)]. The youngest animals included in our study still presented a detectable active thymus at post-mortem but its activity and influence on the disease pattern was not explored.

## CONCLUSIONS

Hamsters and ferrets are valuable animal models to study COVID-19. Hamsters are easy to handle and maintain. Ferrets are more expensive than hamsters and require larger housing space, but a change in their generally active behaviour is a relatively easy clinical sign to detect in comparison with hamsters. The marked transient fatigue reported in the infected ferrets at days 7 and 8 was also reported by others (8,9,11) and likewise is a characteristic of the human COVID-19 condition that could be monitored in preventive or therapeutic experimental studies extending to that time-point. Infected hamsters showed remarkable replication of virus in the lung. In our study, the low challenge dose induced a lung infection very consistent within hamster groups and without severe weight loss seen with higher challenge doses; our protocol is therefore more in line with the 3Rs. Pulmonary infection of ferrets with SARS-Co-V-2 requires higher infectious doses per animal when using the intra tracheal route (8)], and is not systematic (11)]. A higher intranasal infection dose may therefore be required for smaller intra-group variability in future studies. Compared with tissue analysis at post-mortem, longitudinal studies based on the collection (under general anesthesia) of clinical samples such as oral and rectal swabs and nasal washes have the advantage of following the same animal overtime, and therefore have the potential of using fewer animals. Both species provided samples with infectious virus for up to D10, with relevance with excretion and potential of transmission.

## Author Contributions

Conceptualisation; EML, PM, SL and AS; methodology: FB, MW, EPM, SM, SLa and PM; validation: PM, EML, CR, AS and SL; formal analysis: JS, MAG, JV, CP, SM, SLa, EPM, CR and MW; writing—original draft preparation: EML and PM; writing—review and editing, all; figure: Sla; supervision: PM and EML. All authors have read and agreed to the published version of the manuscript.

## Ethical approval

These experiments were approved by the Anses/ENVA/UPEC ethic committee and the French Ministry of Research (Apafis n°24818-2020032710416319).

## Acknowledgments

Euroferret for providing animals very rapidly in times of COVID-19 and severe confinement measures imposed in France. From Anses LRFSN, Jonathan Rieder, Mélanie Badré-Biarnais, Youssef Arnaout, Jean-Marc Boucher, Vanessa Bastid, for their investment in virological and serological analyses ; Valère Brogat, Michel Munier et Nicolas Penel for animal care ; Franca Rizzo for biosecurity and logistical support ; Astrid Vabret for granting access to SARS-CoV2 strains isolated in the University Hospital of Caen, Normandy, France ; Thomas Labadie from London School of Hygiene and Tropical Medicine, UK, for technical advices on viral titration of SARS-CoV2 UCN19 stocks; Latifa Lakhdar head of the Plateforme d’expérimentation animale, Anses - Laboratoire de Lyon

